# Effects of gonadectomy and androgen on neuronal plasticity in motivation and reward related brain regions in the male rat

**DOI:** 10.1101/2020.06.23.166777

**Authors:** Patty T. Huijgens, Eelke M.S. Snoeren, Robert L. Meisel, Paul G. Mermelstein

## Abstract

Gonadal hormones affect neuronal morphology to ultimately regulate behavior. Here, we investigated the effect of both castration and androgen replacement on spine plasticity in the nucleus accumbens shell and core (NAcSh and NAcC), caudate putamen (CPu), medial amygdala (MeA), and medial preoptic nucleus (MPN). Intact and castrated (GDX) male rats were treated with dihydrotestosterone (DHT, 1.5mg) or vehicle (oil) in 3 experimental groups: intact-oil, GDX-oil and GDX-DHT. Spine density and morphology, measured 24 hours after injection, were determined through 3D reconstruction of DiI-labeled dendritic segments. GDX decreased spine density in the MPN, which was rescued by DHT treatment. MeA spine density increased in GDX-DHT animals compared to intact-oil animals. In the NAcSh, DHT decreased spine density, and also rapidly increased the number of pCREB+ cell bodies. These findings indicate that androgen signaling plays a role in the regulation of spine plasticity within neurocircuits involved in motivated behaviors.

## 1. Introduction

Gonadal hormones are key regulators of rewarding behavior (Paredes, 2010; Tonn Eisinger et al., 2018b). Estrogen, progestin, and androgen signaling in the brain is involved in the display of motivated behaviors such as copulation, aggression, and physical activity (Beatty, 1979). Moreover, gonadal hormones have been shown to influence the susceptibility to addiction-like behavior (Tonn Eisinger et al., 2018a). In order to understand how hormones affect behavior, it is important to study the underlying neurobiological mechanisms.

One mechanism through which gonadal hormones exert their influence on motivated behaviors is by affecting the structural plasticity of neurons. Previous research has shown that spine density, spine morphology, and dendrite length can be impacted by gonadal hormones in multiple brain regions involved in motivation (Calizo and Flanagan-Cato, 2002; Cooke and Woolley, 2005; Cunningham et al., 2007; Peterson et al., 2015). These hormone-induced structural reorganizations are both sexually dimorphic and strikingly different between brain regions, and have been linked to motivated behavior, learning, memory, and addiction (Frankfurt and Luine, 2015; McEwen and Milner, 2017; Tobiansky et al., 2018; Tonn Eisinger et al., 2018a).

Copulation is a naturally occurring motivated behavior reliant on gonadal hormones. Earlier research has shown that structural neuronal plasticity could be at the basis of hormonal effects on sexual behavior (Micevych et al., 2017). For example, within the hypothalamus, estradiol appears to enhance neuronal connectivity, essential for lordosis (Meisel and Luttrell, 1990; Dewing et al., 2007; Christensen et al., 2011; Inoue et al., 2019). Estrogens impact additional structures in the female limbic system. For example, spine density in the hippocampus fluctuates during the estrous cycle and estradiol increases spine density in ovariectomized animals (Woolley et al., 1990; Woolley and McEwen, 1992). In contrast, estradiol administration to ovariectomized hamsters or rats produces a decrease in spine density within the NAcC (Staffend et al., 2011; Peterson et al., 2015).

Castration gradually ceases all sexual behavior in male rats and hormonal replacement fully restores copulation (Hull et al., 2006). Yet, in males it remains grossly unknown what neurobiological mechanisms underlie the loss of sexual behavior following loss of gonadal hormones, and whether hormone effects on structural plasticity could be involved. While some studies have shown spine plasticity in response to testosterone in males, it remains unclear to what extent this is mediated by estrogen formed through aromatization of testosterone (Danzer et al., 2001; Chen et al., 2013; Garelick and Swann, 2014). It is, however, evident that estrogen does not simply have the same effects on spine plasticity in males as in females. For example, as mentioned the hippocampal CA1 region exhibits increased spine density upon estrogen treatment in females, but is unresponsive to estrogens in males (Gould et al., 1990; Leranth et al., 2003). Instead, CA1 spine density in males is induced by dihydrotestosterone (DHT), a high-affinity ligand of the androgen receptor that is not aromatized into estradiol (Leranth et al., 2003). Our lab also recently reported similar effects in the nucleus accumbens, where estrogen affects spine plasticity in females and DHT in males, again indicating that these effects in males are caused by androgens rather than estrogens (Peterson et al., 2015; Gross et al., 2018).

Although the effects of gonadal hormones on spine plasticity are sexually dimorphic, there are indications that the underlying mechanisms through which these effects arise are homologous. Specifically, hormone-induced spine plasticity in the nucleus accumbens is mediated by activation of metabotropic glutamate receptor (mGluR) signaling, via estradiol in females and DHT in males (Peterson et al., 2015; Gross et al., 2018). In females, the estrogen-induced spine plasticity is reliant on membrane-bound estrogen receptors that are coupled to mGluRs, which are activated upon estrogen binding to the estrogen receptor. The activation of mGluRs can induce a downstream phosphorylation pathway leading to increased phosphorylation of cAMP response-element binding protein (CREB) (Boulware et al., 2005; Grove-Strawser et al., 2010). CREB is a transcription factor involved in numerous behavioral outputs and implicated in spine plasticity (Lonze and Ginty, 2002; Sargin et al., 2013). Because androgen-induced spine plasticity in the nucleus accumbens in males is also mediated by mGluRs, it could be expected that androgen signaling in males has similar effects on CREB phosphorylation as estradiol in females, perhaps mediated by membrane-associated androgen receptors (Nguyen et al., 2009; Hatanaka et al., 2015; Guo et al., 2020).

In the present study, we investigated the effects of GDX and androgen replacement on neuronal plasticity in putatively important brain regions for sexual motivation in male rats. We hypothesized that GDX could lead to alteration of structural plasticity in the medial preoptic nucleus (MPN), medial amygdala (MeA), and nucleus accumbens core and shell (NAcC and NAcSh), possibly indicating a mechanism for GDX-induced loss of sexual behavior. In addition, we studied how androgen signaling in GDX males impacts structural plasticity. Finally, we built on the hypothesis that the observed effects could be membrane-bound androgen receptor mediated by looking at rapid induction of phosphorylated CREB (pCREB) in the striatum following DHT treatment in GDX males.

## 2. Materials and methods

### 2.1 Animals

Intact and castrated Sprague-Dawley rats (200–225g, 8 weeks old) were purchased from Envigo Laboratories (Indianapolis, IN, United States). Animals were housed 2 per cage (DiI labeling) or 3 per cage (pCREB IHC) and kept on a 12:12 light–dark cycle with food and water *ad libitum*. Animals were allowed to habituate to the research facility for at least 1 week prior to the start of any experiment. All animal procedures were in accordance with the National Institutes of Health Guidelines for the Care and Use of Laboratory Animals and were approved by the Animal Care and Use Committee at the University of Minnesota.

### 2.2 Treatment, tissue processing and DiI labeling

5α-androstan-17β-ol-3-one (DHT; Steraloids Inc.; Newport, RI, United States) was dissolved in cottonseed oil. Ten to 30 days after arrival the rats were injected s.c. with 1.5 mg DHT or vehicle in a volume of 0.2 mL. The experiment was run in batches of 2 animals (cage mates) at a time. The two animals in each batch were in the same group, and treatment groups were randomized according to a latin square design, so that average castration duration did not differ between castrated groups. Animals were sacrificed 24 hours after hormone or vehicle treatment.

The tissue was prepared and ballistically labeled according to protocol as described previously (Staffend and Meisel, 2011). Animals were euthanized by Euthasol overdose (0.35 mL i.p; 390 mg/mL pentobarbital sodium, 50 mg/mL phenytoin sodium, Virbac AH Inc., Nice, France), injected with 0.25 mL heparin into the left ventricle, and transcardially perfused with 50 mL 25 mM phosphate-buffered saline (PBS, pH = 7.2) followed by 500 mL 1.5% paraformaldehyde in PBS. Brains were removed and post-fixed in 1.5% paraformaldehyde for 1 hour. Then, brains were sliced coronally into 300 µm thick sections using a Leica VT1000 S Vibratome (Buffalo Grove, IL, United States). Sections containing the brain regions of interest, i.e. the caudate putamen (CPu), nucleus accumbens core and shell (NAcC and NAcSh), medial preoptic nucleus (MPN) and medial amygdala (MeA), were collected and stored in PBS until ballistic labeling

DiI bullets were prepared from Tefzel tubing (Bio-Rad, Hercules, CA, United States) pretreated with 15 mg/ml polyvinylpyrrolidone (PVP) in deionized water. Two milligrams of DiI (Molecular Probes, Carlsbad, CA, United States) was dissolved in 100 μl dichloromethane and applied to 90 mg of 1.3 μm tungsten microcarrier particles (Bio-Rad) spread out evenly on a glass slide. The coated tungsten particles were suspended in 10 mL PVP solution, and disaggregated by sonication and intermittent vortexing for 12 min. The pretreated Tefzel tubing was subsequently coated with the DiI-tungsten particles by allowing the suspension to settle in the tubing for 3 min, after which the suspension was quickly expelled. The tubing was dried under 0.4 LPM nitrogen gas flow using a tubing prep station (Bio-Rad) for 30 min, after which the tubing was cut into 1.3 cm long “bullets”. Bullets were loaded into a Helios Gene Gun (modified barrel, 40 mm spacer, 70 μm mesh filter; Bio-Rad) and PBS surrounding brain sections was removed. DiI-tungsten particles were shot into the tissue by shooting 1 bullet on each section using helium gas expulsion (100 PSI). To allow DiI spreading throughout the labeled neurons, sections were kept overnight in PBS in the dark. The next day, sections were postfixed in 4% paraformaldehyde for 1 hour, rinsed in PBS, mounted on slides and coverslipped with FluorGlo mounting media for lipophilic dyes (Spectra Services, Ontario, NY, United States). Note that the FluoroGlo mounting medium is no longer available.

### 2.3 DiI confocal imaging, reconstruction and quantitation

Using a Leica TCS SPE confocal microscope, brain regions of interest were identified and delineated using low magnification brightfield in conjunction with the Paxinos & Watson rat brain atlas (6^th^ edition) for reference. For each brain region, 2-3 dendritic segments (70-200 µm away from soma, and more than 10 µm away from dendritic end points and bifurcations) per neuron, in 2-3 neurons, were imaged and analyzed (Figure 1). Dendritic segments were imaged using a Leica PLAN APO 63×, 1.4 NA oil immersion objective (11506187, Leica, Mannheim, Germany) and Type LDF immersion oil (Cargille, Cedar Grove, NJ, United States). All images were taken at an *xy* pixel distribution of 512 × 512, a frequency of 400 Hz, a step size of 0.12 μm and optical zoom of 5.6, with the laser power and photomultiplier being adjusted to capture the dendrite in its full dynamic range. Data from 9 to 10 animals were collected for each treatment group. In case there were less than 2 neurons in a brain region feasible for imaging, the animal was excluded from further analysis for that region. This predominantly occurred in the MPN and MeA, and explains the smaller sample sizes for these regions.

**Figure 1.**
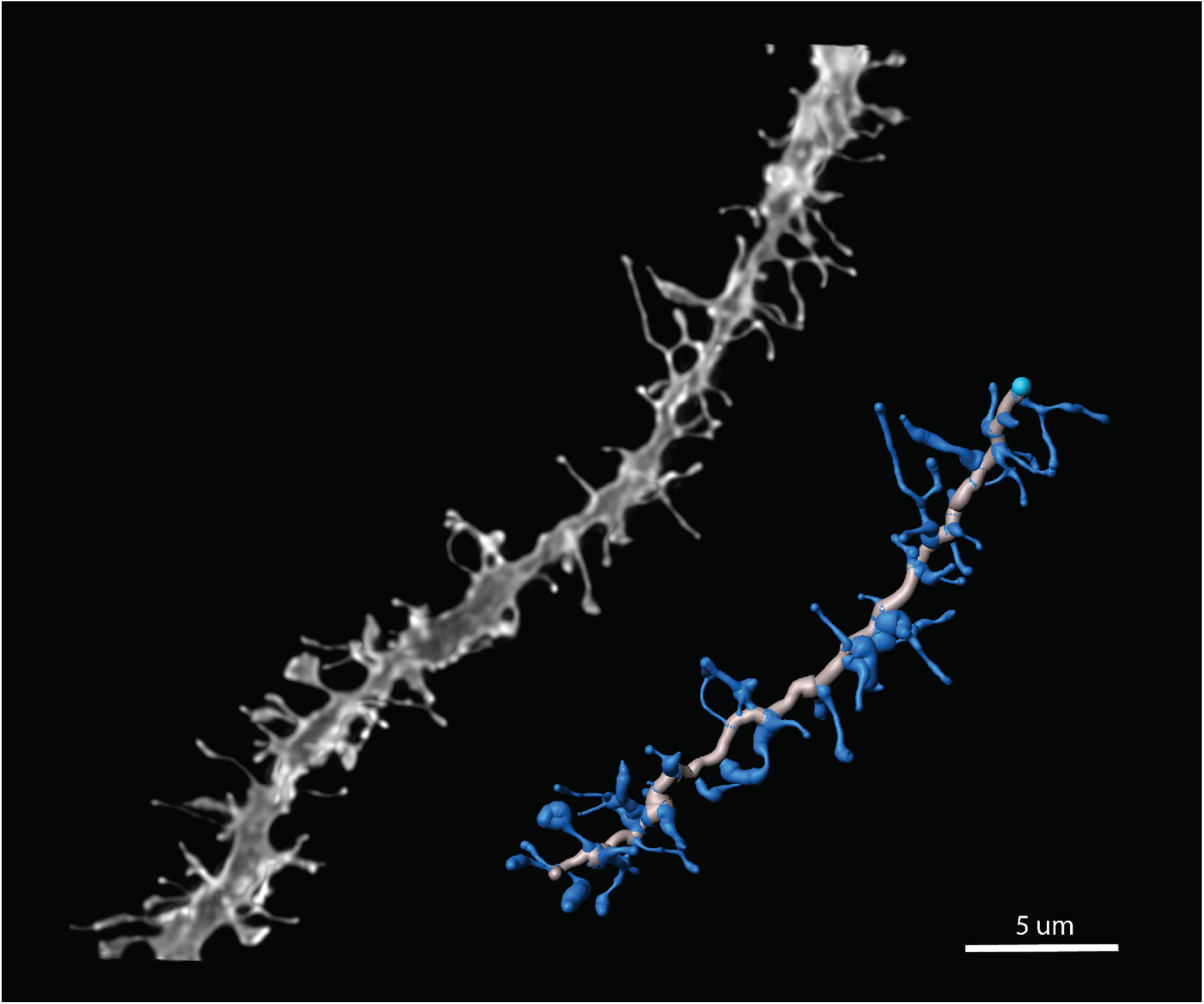
Dendritic segment reconstruction. Representative maximum projection image and 3D reconstruction process of a striatal medium spiny neuron dendritic segment labeled with DiI. Image acquired at 63x, scale bar 5 µm.

After imaging, optical sections were processed through 3D deconvolution using AutoQuant X3 AutoDeblur software (Media Cybernetics, Bethesda, MD, United States). Deconvoluted z-stacks were then reconstructed in the Surpass module of Imaris software (Bitplane Inc., Concord, MA, United States), through manual tracing of dendrites and spines using the Filament tool and Autodepth function. A 3D reconstruction of 15-20 µm of dendritic shaft and spines was rendered using the diameter function with a contrast threshold of 0.7, and data on spine density, spine length, and head diameter was collected for each segment. Spine densities for each segment (collected as average spine density/10 μm) were averaged across each neuron and then within each brain region for each animal, providing a region-specific spine density average for each animal that was then used for statistical analysis. Measurements of spine length and head diameter were pooled for each treatment condition and then plotted in violin plots, as well as binned cumulative probability distributions (bin sizes: spine length, 0.5 μm; head diameter, 0.1 μm).

### 2.4 pCREB immunohistochemistry

2-hydroxypropyl-β-cyclodextrin (cyclodextrin; Sigma-Aldrich) was dissolved in sterile water to obtain a 45% vehicle solution. DHT was dissolved in cyclodextrin solution, and 1.5 mg (high dose) or 0.15 mg (low dose) was injected i.p. in a volume of 0.2 mL. After injection, animals were put alone in a cage until lethal i.p. injection with Euthasol (0.35 mL i.p; 390 mg/mL pentobarbital sodium, 50 mg/mL phenytoin sodium, Virbac AH Inc.) 15 or 30 min later. Then, animals were injected with 0.25 mL heparin into the left ventricle, and transcardially perfused with 50 mL PBS followed by 500 mL 4% paraformaldehyde in PBS. Brains were removed and post-fixed in 4% paraformaldehyde for 2 hours, and stored in 10% sucrose in PBS overnight at 4°C. The next day, brains were cut on a freezing microtome into 40 µm sections and every third section throughout the striatum was collected into 0.1% BSA in 25 mM PBS (BSA/PBS) for immediate free-floating immunohistochemical processing. After rinsing in BSA/PBS, sections were incubated in polyclonal rabbit anti-phospho-CREB (1:2000; cat. 06-519, Merck Millipore) in 0.3% Triton-X-100 in BSA/PBS for 48 hours at 4°C. Subsequently, sections were incubated in biotylinated goat anti-rabbit (1:200; VECTASTAIN Elite ABC-HRP rabbit-IgG Kit, Vector Laboratories) in BSA/PBS for 1 hour, avidin-biotin-peroxidase complex (1:100; VECTASTAIN Elite Kit) in PBS for 1 hour, and 3,3’-diaminobenzidine (0.8 mg/mL; Sigma-Aldrich) with 0.3% H_2_O_2_ in 50 mM Tris buffer (pH 7.6) for 8 min, with repeated buffer washing in between all steps. Sections were then mounted on slides, and coverslipped using DPX mounting medium (Sigma-Aldrich). The experiment was run in batches of 3 animals at a time, with the same treatment injection timing (15 or 30 min) for each animal in the batch, and 1 animal per treatment group per batch. Administration of different treatments was randomized according to a latin square design so that the order of injection and perfusion would not be a factor.

### 2.5 pCREB imaging and quantitation

For each animal, 3 sections within the central striatum were identified and imaged using a Leica DM 4000 B LED microscope and 10x objective. At the level of the nucleus accumbens the anterior commissure has a lateral monotonic migration. Consequently, sections were matched on the anterior-posterior axis by selecting those sections in which the distance from the tip of the lateral ventricle to the medial edge of the anterior commissure was 300 - 350 µm. Images were always taken on the right side of the section, without avoidance of artifacts. The same exposure and white balance settings were used across all images. The images were subsequently loaded in Adobe Photoshop and a red box (300 × 500 µm) placed within the brain regions of interest. For the NAc, the box was placed medial to the ventricle for the shell and lateral to the ventricle for the core, and the distance between the boxes was kept at 100 µm for each section imaged. For the medial and lateral CPu, the top corner of the boxes touched the corpus callosum.

For pCREB+ cell counting, the cell counter plugin in ImageJ software was used. In order to increase intra-observer reliability, we converted the images to greyscale and used the automated threshold algorithm “Otsu” to acquire a binary image that separated positive cells from background to use as a counting guide. Otsu’s method finds a threshold value where foreground and background pixel value variance is at a minimum. Because some batch-to-batch immunohistochemistry variance is to be expected, Otsu’s method works well for thresholding here because it uses information from within the image to separate background from staining. Cells were counted if they appeared with at least 1 pixel in the thresholded image, and standard stereology rules were applied when counting on the box borders. Intra-observer agreement of positive cell count was within 97.4-99.5% for a sample size of 10 duplicate images. Cell counts of 3 sections were averaged across each brain region within each animal. Out of 384 total boxes, 8 images contained very large artefacts in the tissue within the box, and were therefore excluded from analysis.

### 2.6 Data analysis and statistics

All data analysis was conducted in GraphPad Prism 8 (GraphPad Software, San Diego, California USA). For spine density and pCREB expression, groups were compared using a one-way ANOVA, followed by Tukey’s multiple comparisons test in case of significant effect. The binned spine morphology probability distributions were compared to each other group using a Kolmogorov-Smirnov test.

## 3. Results

### 3.1 Dendritic spine plasticity

To study the effects of both GDX and androgen replacement on dendritic spine plasticity, we compared intact males treated with oil (vehicle) to GDX males treated with oil, to GDX males treated with DHT. We found that GDX affected spine plasticity in the MPN (Figure 2A, [F_(2,15)_=10.70, p=0.0013, η^2^ = 0.59]) by decreasing spine density [mean diff. vs. intact = 2.03, p=0.0037, *g* = 2.35]. GDX did not affect spine density in the MeA, NAcSh, NAcC, and CPu (Figure 2A). DHT administration to GDX animals rescued the GDX-induced spine loss in the MPN [mean diff. vs. GDX-oil = 2.15, p=0.0036, *g* = 2.54]. In contrast, DHT decreased spine density in the NAcSh in GDX animals compared to oil-treated GDX animals (Figure 2A, [F_(2,25)_=3.56, p=0.04369, η^2^ = 0.22], [mean diff. vs. GDX-oil = 2.54, p=0.0341, *g* = 1.44]). This effect, however, was not different compared to the intact-oil group. We found effects on spine density in the MeA as well (Figure 2A, [F_(2,21)_=4.45, p=0.0245, η^2^ = 0.30]). Specifically, although GDX itself did not affect spine density, DHT treated GDX males had a higher spine density than oil treated intact males [mean diff. vs. intact = 2.71, p=0.0193, *g* = 1.56]. We saw no effects of DHT on spine density in NAcC and CPu (Figure 2A), neither compared to the intact-oil nor the GDX-oil group.

**Figure 2.**
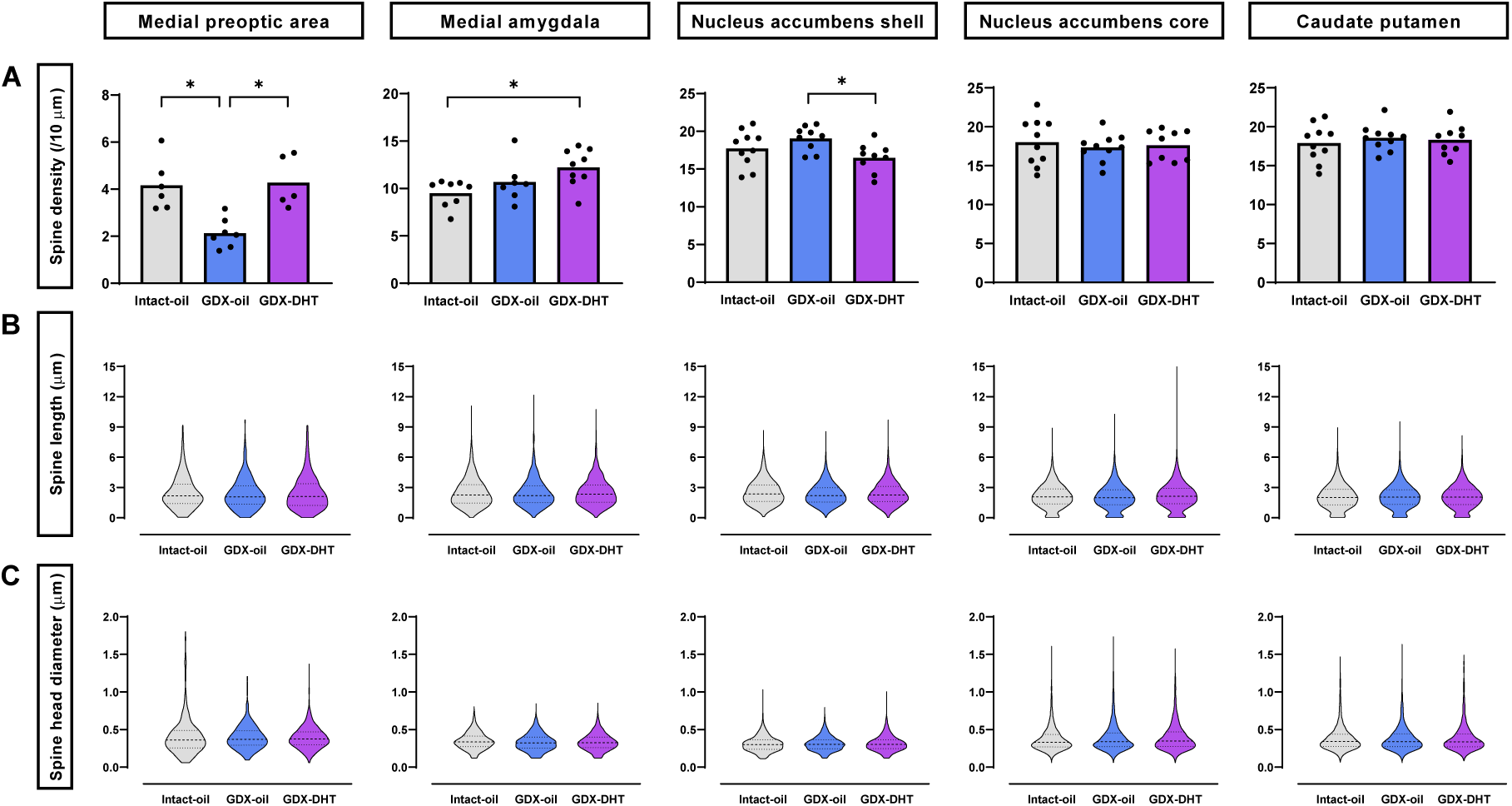
Gonadectomy (GDX) and dihydrotestosterone (DHT) affect spine plasticity differentially across several brain regions regulating motivated behavior. **(A)** Spine density 24 hours after treatment with oil or DHT in intact and GDX males in the medial preoptic area (MPN), medial amygdala (MeA), nucleus accumbens shell and core (NAcSh and NAcC) and caudate putamen (CPu). Individual values represent neuron spine density average per animal (= unit of analysis), which is comprised of the average spine density across 2-3 neurons per animal, calculated from the average spine density from 2-3 segments per neuron. n = 6, 7, 5 (MPN); 8, 7, 9 (MeA); 10, 10, 9 (NAcC); 10, 9, 9 (NAcSh); 10, 10, 9 (CPu) animals per group. *p<0.05. **(B)** Violin plot representation of spine length distribution. For violin plots, all spine data points from all animals within the same group were pooled into one plot. Dashed line, median; dotted lines, quartiles. **(C)** Violin plots of spine head diameter distribution.

Spine morphology gives information about spine maturation and function (Rochefort and Konnerth, 2012). Because spine morphology is determined by spine length and spine head diameter, we compared the distributions of these two parameters between groups. No effects of castration or DHT treatment were observed on spine length or spine head diameter in any of the brain regions (Figure 2B-C).

### 3.2 pCREB expression

To study the potential rapid effects of DHT we focused on the striatum, a brain region in which rapid phosphorylation of CREB has been documented (Grove-Strawser et al., 2010). We determined the number of cells expressing pCREB by means of immunohistochemistry (Figure 3A), 15 and 30 min after hormone or vehicle treatment of GDX males. DHT treatment had no effect on the amount of pCREB+ cells within 15 minutes of hormone administration in any of the investigated subregions (Figure 3B). After 30 min, however, a high dose of DHT significantly increased the number of pCREB+ cells in the NAcSh (Figure 3C [F_(2,12)_=5.039, p=0.0258, η^2^ = 0.46]) compared to vehicle [mean diff. 90, p=0.0358, *g* = 1.79]. A low dose of DHT also increased the number of pCREB+ cells in the medial CPu (Figure 3C [F_(2,12)_=4.350, p=0.038, η^2^ = 0.42]) compared to vehicle [mean diff. 82, p=0.0321, *g* = 1.77]. No effects of DHT treatment were found in the NAcC and lateral part of the CPu.

**Figure 3.**
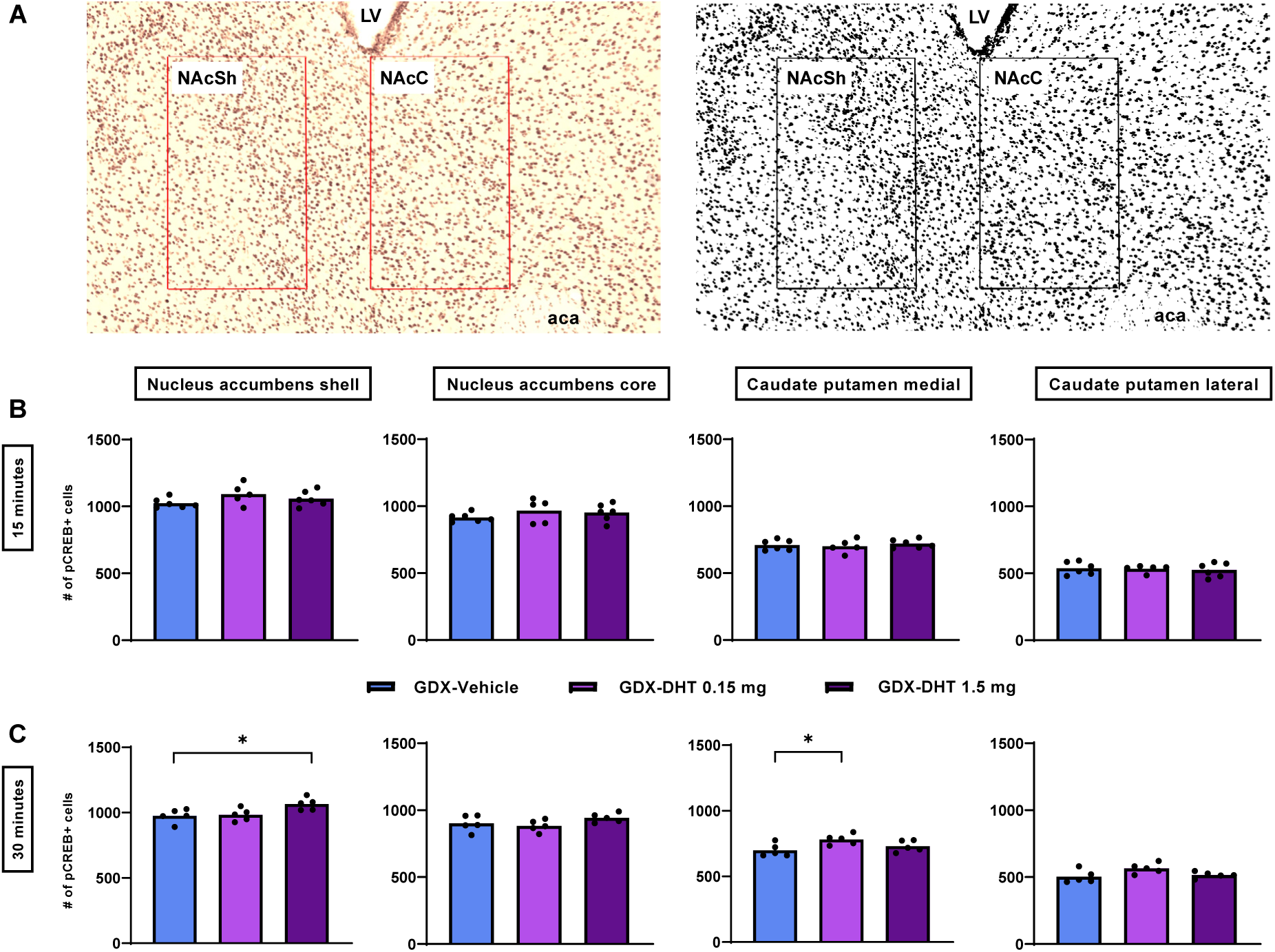
DHT rapidly induces CREB phosphorylation. **(A)** Representative image of pCREB staining and counting box (300 µm x 500 µm) delineating in the nucleus accumbens core and shell, and corresponding Otsu thresholded image. Magnification is illustrated by the boxes outlining the region of interest (300 µm X 500 µm). LV = lateral ventricle, aca = anterior commissure. **(B)** Number of pCREB positive cells in each brain region 15 min after i.p. treatment with oil or low or high dose DHT in GDX males. **(C)** Number of pCREB positive cells 30 min after treatment. *p<0.05.

## 4. Discussion

Gonadal hormones are known to regulate synaptic plasticity (Parducz et al., 2006; Hyer et al., 2018). Whereas the literature has so far mostly characterized the effects of estrogen in females, some evidence exists for effects of gonadal steroids in males as well (Leranth et al., 2003; Hajszan et al., 2007; Hatanaka et al., 2015). With the current study, we aimed to examine the effects of loss and subsequent replacement of gonadal hormones on spine plasticity in males. We focused on brain regions that are involved in neural circuits of (sexual) motivation: the NAcC, NAcSh and CPu, which are part of dopaminergic reward processing, and the MPN and MeA, two regions critically involved in the display of sexual behavior. The studied brain regions have all been shown to be androgen target areas in rats (Simerly et al., 1990). We showed that GDX decreased spine density in the MPN, which was rescued by DHT treatment. In addition, DHT decreased spine density in the NAcSh in GDX animals, whereas it increased spine density in the MeA of GDX animals compared to intact animals. Thus, spine plasticity is differentially affected by gonadal hormones across the studied brain regions.

Gonadectomy gradually ceases copulation in male rats as a result of the loss of gonadal hormones. Estrogen as well as androgen signaling through estrogen and androgen receptors in the brain is necessary for the full display of male sexual behavior (Hull et al., 2006). Considering the high expression of androgen and estrogen receptors in the MPN, it is therefore not surprising that the MPN is the most important brain region for regulation of sexual behavior in males (Hull et al., 2006). Disruption of the MPN through lesions causes gonadally intact male rats to stop copulating (Hull et al., 2006). Local infusion of an aromatase inhibitor (preventing the formation of estradiol and thus estrogen receptor signaling) or an androgen antagonist (preventing androgen receptor signaling) into the MPN, both suppress copulation in gonadally intact male rats, showing a vital interaction of gonadal hormones and the MPN in male sexual behavior (Clancy et al., 1995; McGinnis et al., 2002). Yet, it remains unclear what mechanism underlies the importance of the activity of gonadal hormones in the MPN for copulation in male rats. What *has* been shown earlier is that GDX reduces dopamine release and c-Fos expression in the male’s MPN upon exposure to an estrous female (Hull et al., 1995; Paredes et al., 1998; Putnam et al., 2003). This suggests that a lack of gonadal hormones may reduce afferent sensory information to the MPN. With our current study, we demonstrate a novel effect of GDX in the MPN of male rats. GDX drastically reduces spine density of MPN neurons, an indication of an overall decrease in synapses within the MPN. In line with reduced dopamine release and c-Fos expression in the MPN, this suggests a model in which gonadal hormones act as facilitators contributing to MPN connectivity. This connectivity may then be necessary for sexual behavior to arise in response to the stimulus of an estrous female. Our study shows that the GDX-induced spine loss is present in males gonadectomized for longer than 10 days, a time point at which most male rats would have stopped copulating (Hull et al., 2006). Future research should focus on spine plasticity at different time points after GDX in order to reveal whether loss of spines in the MPN also coincides with the gradual loss of sexual behavior after GDX in males. That should provide more insight in whether GDX-induced loss of spines in the MPN indeed contributes to loss of sexual behavior.

Treatment with testosterone given systemically or locally into the MPN facilitates copulation in GDX males (McDonald et al., 1970; Feder, 1971; Johnston and Davidson, 1972; McGinnis and Dreifuss, 1989). In addition, testosterone rescues GDX induced spine loss in the MPN of male hamsters (Garelick and Swann, 2014). Furthermore, functional aromatization of testosterone into estrogen is necessary for the display of the full range of sexual behavior in male rats. Treatment of GDX males with DHT, a high-affinity ligand of the androgen receptor that cannot be aromatized into estrogen, is ineffective in reinstating sexual behavior (Johnston and Davidson, 1972). Further, at the same time that DHT has affinity for estrogen receptor β (ERβ), our prior work showed that an ERβ agonist did not affect dendritic spine density in the nucleus accumbens (Gross et al., 2018). Therefore, we expected to find that treatment with DHT would not be sufficient to rescue the GDX-induced spine loss in the MPN. Nevertheless, we show here that DHT treatment of GDX males *does* fully restore spine density on MPN neurons. Even though DHT-induced spinogenesis in the MPN of GDX males does not coincide with restoration of copulation (Christensen and Clemens, 1974), androgen signaling still contributes to copulatory behavior. For example, Local infusion of an androgen receptor antagonist into the MPN of GDX males prevents the reinstatement of sexual behavior by systemic testosterone treatment (Harding and McGinnis, 2004). In line with this, androgen signaling in addition to estrogen signaling is necessary for the motivational aspects of sexual behavior such as preference for an estrous female, and DHT alone has mild effects on sexual incentive motivation (Matuszczyk and Larsson, 1994; Attila et al., 2010). Furthermore, testosterone and estradiol both show rapid effects on firing rate in MPN neurons, but they rarely affect the same neurons (Silva and Boulant, 1986). Therefore, androgenic signaling may perhaps primarily influence and maintain sexual motivation through a distinct neuronal population in the MPN, mediated by spine plasticity. An important research focus in the future will be to unravel the effects of estrogen on spine plasticity in the MPN of GDX males.

In contrast to our findings in the MPN, we found that GDX did not decrease spine density in the MeA, another important region for copulation (Hull et al., 2006). Others have reported a decrease in spine density in the posterodorsal MeA, 3 months after GDX, measured on dendrites very proximal to the soma (de Castilhos et al., 2008; Zancan et al., 2017), and in males castrated before puberty (Cooke and Woolley, 2009). Our measures, on the other hand, were taken at least 70 µm away from the soma, not on primary dendrites, and within a shorter time frame after GDX. We did find, however, that DHT has spinogenic properties in the MeA, even though we only saw this in comparison to intact males. Possibly, gonadal hormones are not necessary to maintain spine density in the MeA, but do have the ability to affect spine plasticity like in the MPN. Another study conducted in intact pubertal males showed that a chronic high dose testosterone is transiently spinogenic in the antero- and posterodorsal MeA (Cunningham et al., 2007). Thus, gonadal hormones may affect MeA spine plasticity differentially depending on the distance of a dendritic segment from the soma, the timing of castration within life, the castration duration, and the amount of time that has gone by since hormone treatment.

The current experiment replicated earlier results from our lab, where we showed that in contrast to its spinogenic effects in the MPN and MeA, DHT decreases spine density after GDX in NAcSh, but not in NAcC and CPu in GDX males (Gross et al., 2018). Here we used an additional control group of intact males to also establish that gonadal hormones are not necessary to maintain spine density and morphology in the striatum in males, as GDX left these variables unaffected. Another group found that in intact males, a chronic high dose of testosterone decreases spine density in the NAcSh, and has no effect on the NAcC (Wallin-Miller et al., 2016). This suggests that androgens induce loss of spines in the NAcSh regardless of whether the male is gonadectomized or not. The rapid changes in NAcSh dendritic spines following DHT do not appear to underlie the expression of copulation in males as the DHT effects on spines requires mGluR5 signaling (Gross et al., 2018) and accumbens antagonism of mGluR5 receptors does not disrupt copulation (Pitchers et al., 2016). The medial preoptic area and medial amygdala are better candidates for sites of action of DHT on copulation, and DHT modulated dendritic spines within 24 hours in these regions as well. One study has demonstrated interactions between estrogen receptor and mGluR signaling in the medial preoptic area (Santollo and Daniels, 2019) with nothing known about similar interactions in the medial amygdala. As both estradiol and DHT induce rapid ERK phosphorylation in the medial preoptic area (Jean et al., 2017), cooperative signaling through mGluR receptors could be the basis for rapid effects on copulation in males. We propose parallels with the mechanisms through which estradiol acts to regulate female sexual behavior. Estrogens induce rapid membrane-mediated signaling cascades, which are followed by longer lasting transcriptional activation via nuclear receptors (Vasudevan et al., 2005). We envision a similar set of actions for male sexual behavior in which androgens provide both rapid and long-term plasticity.

The small numbers of dendritic spines measured in these studies sometimes raise questions about the functional significance of these spine changes. For striatal medium spiny neurons, Golgi studies suggest that the cumulative dendritic length may be on the order of 2100 µm (Bicanic et al., 2017), while cell fills put the number closer to about 3000 µm (Gertler et al., 2008). With an increase of 3 spines/10 µm as we see in the nucleus accumbens shell, this translates to upwards of 1000 excitatory synapses per medium spiny neuron, producing a substantial impact on the electrotonic potential of these neurons (Gertler et al., 2008).

Androgens can exert their action on neurons through multiple signaling pathways (Foradori et al., 2008). While the 24 hours after hormone treatment in this experiment is long enough for genomic effects to occur, previous results from our lab show that DHT-induced spine plasticity in the NAcSh is mediated by mGluR5, a G protein-coupled receptor associated with the Gαq protein, suggesting that membrane-initiated signaling pathways are involved (Gross et al., 2018). This mechanism is homologous to the mGluR5 mediated estrogen-induced decrease in spine density in the NAcC of ovariectomized females (Peterson et al., 2015). The coupling to and regulation of mGluRs by membrane-bound estrogen receptors has been well characterized and has been shown to mediate spine plasticity and behavior in females (Micevych and Mermelstein, 2008; Christensen et al., 2011; Boulware et al., 2013). Estrogen rapidly increases phosphorylation of the transcription factor CREB through its membrane interaction with mGluRs in hippocampal and striatal neurons exclusively in female cultures (Boulware et al., 2005; Grove-Strawser et al., 2010). It is important to note that while estrogen receptor/mGluR signaling leads to pCREB across many brain regions, only a subset of these exhibit changes in dendritic spines. Therefore, this signaling pathway may be necessary for structural changes, but it is not sufficient. In male cultures, estradiol does not induce pCREB, while mGluR activation does. In addition, activation of mGluR5 mediates spine plasticity in male NAcC and NAcSh (Gross et al., 2016), suggesting that mGluRs possibly couple to membrane bound androgen receptor in males. The androgen receptor has indeed also been shown to migrate to the membrane (Tabori et al., 2005), using the same intracellular processes as estrogen receptors (Pedram et al., 2007). Here, we show that DHT is capable of inducing striatal pCREB in vivo within 30 min of injection. While the immunohistochemistry method that we used for assessing pCREB expression only allowed for counting of number of positive cells, and not the level of pCREB within a cell, it is possible that DHT *also* induces higher phosphorylation levels of pCREB in each individual positive cell. Still, our results point towards a pathway in which androgen binds to membrane-bound androgen receptors, which activates mGluR5 though coupling in the NAcSh but not in the NAcC. This leads to activation of a downstream signaling cascade culminating into phosphorylation of CREB, thereby enhancing its gene transcription properties. There is a large body of literature on the function of CREB, which among others is involved in learning and memory and synaptic plasticity (Lonze and Ginty, 2002). Whether this proposed mechanism of DHT-induced plasticity, through mGluR5 or other mGluRs, can also be applied to the effects we found in the MPN and MeA will be part of future research.

## 5. Conclusion

We conclude that both GDX and androgen differentially affect spine plasticity in the MPN, MeA, and NAcSh, while NAcC and CPu remain unaffected. In the NAcSh, DHT may exert its effects through pCREB induction mediated by androgen receptor activation of mGluR5.

## Acknowledgements

Financial support was received from NIH DA041808 to PGM and from Norwegian Research Council grant #251320 to EMS. PTH was supported by a Personal Overseas Research Grant from the Norwegian Research Council. We thank Anna Peyla for technical assistance with immunohistochemistry.

Declarations of interest: none

## References

Attila, M., Oksala, R., and Ågmo, A. (2010). Sexual incentive motivation in male rats requires both androgens and estrogens. Horm Behavr 58, 341–351. doi: 10.1016/j.yhbeh.2009.08.011.

Beatty, W.W. (1979). Gonadal hormones and sex differences in nonreproductive behaviors in rodents: organizational and activational influences. Horm Behav 12, 112–163. doi: 10.1016/0018-506x(79)90017-5.

Bicanic, I., Hladnik, A., and Petanjek, Z. (2017). A quantitative Golgi study of dendritic morphology in the mice striatal medium spiny neurons. Front Neuroanat 11, 37. doi: 10.3389/fnana.2017.00037.

Boulware, M.I., Heisler, J.D., and Frick, K.M. (2013). The memory-enhancing effects of hippocampal estrogen receptor activation involve metabotropic glutamate receptor signaling. J Neurosci 33, 15184–15194. doi: 10.1523/jneurosci.1716-13.2013.

Boulware, M.I., Weick, J.P., Becklund, B.R., Kuo, S.P., Groth, R.D., and Mermelstein, P.G. (2005). Estradiol activates group I and II metabotropic glutamate receptor signaling, leading to opposing influences on cAMP response element-binding protein. J Neurosci 25, 5066–5078. doi: 10.1523/jneurosci.1427-05.2005.

Calizo, L.H., and Flanagan-Cato, L.M. (2002). Estrogen-induced dendritic spine elimination on female rat ventromedial hypothalamic neurons that project to the periaqueductal gray. J Comp Neurol 447, 234–248. doi: 10.1002/cne.10223.

Chen, J.R., Wang, T.J., Lim, S.H., Wang, Y.J., and Tseng, G.F. (2013). Testosterone modulation of dendritic spines of somatosensory cortical pyramidal neurons. Brain Struct Funct 218, 1407–1417. doi: 10.1007/s00429-012-0465-7.

Christensen, A., Dewing, P., and Micevych, P. (2011). Membrane-initiated estradiol signaling induces spinogenesis required for female sexual receptivity. J Neurosci 31, 17583–17589. doi: 10.1523/jneurosci.3030-11.2011.

Christensen, L.W., and Clemens, L.G. (1974). Intrahypothalamic implants of testosterone or estradiol and resumption of masculine sexual behavior in long-term castrated male rats. Endocrinology 95, 984–990. doi: 10.1210/endo-95-4-984.

Clancy, A.N., Zumpe, D., and Michael, R.P. (1995). Intracerebral infusion of an aromatase inhibitor, sexual behavior and brain estrogen receptor-like immunoreactivity in intact male rats. Neuroendocrinology 61, 98–111. doi: 10.1159/000126830.

Cooke, B.M., and Woolley, C.S. (2005). Gonadal hormone modulation of dendrites in the mammalian CNS. J Neurobiol 64, 34–46. doi: 10.1002/neu.20143.

Cooke, B.M., and Woolley, C.S. (2009). Effects of prepubertal gonadectomy on a male-typical behavior and excitatory synaptic transmission in the amygdala. Dev Neurobiol 69, 141–152. doi: 10.1002/dneu.20688.

Cunningham, R.L., Claiborne, B.J., and McGinnis, M.Y. (2007). Pubertal exposure to anabolic androgenic steroids increases spine densities on neurons in the limbic system of male rats. Neuroscience 150, 609–615. doi: 10.1016/j.neuroscience.2007.09.038.

Danzer, S.C., McMullen, N.T., and Rance, N.E. (2001). Testosterone modulates the dendritic architecture of arcuate neuroendocrine neurons in adult male rats. Brain Res 890, 78–85. doi: 10.1016/s0006-8993(00)03083-3.

de Castilhos, J., Forti, C.D., Achaval, M., and Rasia-Filho, A.A. (2008). Dendritic spine density of posterodorsal medial amygdala neurons can be affected by gonadectomy and sex steroid manipulations in adult rats: a Golgi study. Brain Res 1240, 73–81. doi: 10.1016/j.brainres.2008.09.002.

Dewing, P., Boulware, M.I., Sinchak, K., Christensen, A., Mermelstein, P.G., and Micevych, P. (2007). Membrane estrogen receptor-alpha interactions with metabotropic glutamate receptor 1a modulate female sexual receptivity in rats. J Neurosci 27, 9294–9300. doi: 10.1523/jneurosci.0592-07.2007.

Feder, H.H. (1971). The comparative actions of testosterone propionate and 5 -androstan-17 -ol-3-one propionate on the reproductive behaviour, physiology and morphology of male rats. J Endocrinol 51, 241–252. doi: 10.1677/joe.0.0510241.

Foradori, C.D., Weiser, M.J., and Handa, R.J. (2008). Non-genomic actions of androgens. Front Neuroendocrinol 29, 169–181. doi: 10.1016/j.yfrne.2007.10.005.

Frankfurt, M., and Luine, V. (2015). The evolving role of dendritic spines and memory: Interaction(s) with estradiol. Horm Behav 74, 28–36. doi: https://doi.org/10.1016/j.yhbeh.2015.05.004.

Garelick, T., and Swann, J. (2014). Testosterone regulates the density of dendritic spines in the male preoptic area. Horm Behav 65, 249–253. doi: 10.1016/j.yhbeh.2014.01.008.

Gertler, T.S., Chan, C.S., and Surmeier, D.J. (2008). Dichotomous anatomical properties of adult striatal medium spiny neurons. J Neurosci 28, 10814–10824. doi: 10.1523/jneurosci.2660-08.2008.

Gould, E., Woolley, C.S., Frankfurt, M., and McEwen, B.S. (1990). Gonadal steroids regulate dendritic spine density in hippocampal pyramidal cells in adulthood. J Neurosci 10, 1286–1291. doi: 10.1523/jneurosci.10-04-01286.1990.

Gross, K.S., Brandner, D.D., Martinez, L.A., Olive, M.F., Meisel, R.L., and Mermelstein, P.G. (2016). Opposite effects of mGluR1a and mGluR5 activation on nucleus accumbens medium spiny neuron dendritic spine density. PLoS One 11, e0162755. doi: 10.1371/journal.pone.0162755.

Gross, K.S., Moore, K.M., Meisel, R.L., and Mermelstein, P.G. (2018). mGluR5 mediates dihydrotestosterone-induced nucleus accumbens structural plasticity, but not conditioned reward. Front Neurosci 12, 855. doi: 10.3389/fnins.2018.00855.

Grove-Strawser, D., Boulware, M.I., and Mermelstein, P.G. (2010). Membrane estrogen receptors activate the metabotropic glutamate receptors mGluR5 and mGluR3 to bidirectionally regulate CREB phosphorylation in female rat striatal neurons. Neuroscience 170, 1045–1055. doi: 10.1016/j.neuroscience.2010.08.012.

Guo, G., Kang, L., Geng, D., Han, S., Li, S., Du, J., et al. (2020). Testosterone modulates structural synaptic plasticity of primary cultured hippocampal neurons through ERK - CREB signalling pathways. Mol Cell Endocrinol 503, 110671. doi: 10.1016/j.mce.2019.110671.

Hajszan, T., MacLusky, N.J., Johansen, J.A., Jordan, C.L., and Leranth, C. (2007). Effects of androgens and estradiol on spine synapse formation in the prefrontal cortex of normal and testicular feminization mutant male rats. Endocrinology 148, 1963–1967. doi: 10.1210/en.2006-1626.

Harding, S.M., and McGinnis, M.Y. (2004). Androgen receptor blockade in the MPOA or VMN: effects on male sociosexual behaviors. Physiol Behav 81, 671–680. doi: 10.1016/j.physbeh.2004.03.008.

Hatanaka, Y., Hojo, Y., Mukai, H., Murakami, G., Komatsuzaki, Y., Kim, J., et al. (2015). Rapid increase of spines by dihydrotestosterone and testosterone in hippocampal neurons: Dependence on synaptic androgen receptor and kinase networks. Brain Res 1621, 121–132. doi: 10.1016/j.brainres.2014.12.011.

Hull, E.M., Du, J., Lorrain, D.S., and Matuszewich, L. (1995). Extracellular dopamine in the medial preoptic area: implications for sexual motivation and hormonal control of copulation. J Neurosci 15, 7465–7471. doi: 10.1523/jneurosci.15-11-07465.1995.

Hull, E.M., Wood, R.I., and Mckenna, K.E. (2006). “The neurobiology of male sexual behavior. In: The Physiology of Reproduction, third ed. J. Neill, Editor in Chief, Donald Pfaff, Section Editor, Elsevier Press, pp. 1729–1824.

Hyer, M.M., Phillips, L.L., and Neigh, G.N. (2018). Sex differences in synaptic plasticity: Hormones and beyond. Front Molec Neurosci 11. doi: 10.3389/fnmol.2018.00266.

Inoue, S., Yang, R., Tantry, A., Davis, C.H., Yang, T., Knoedler, J.R., et al. (2019). Periodic remodeling in a neural circuit governs timing of female sexual behavior. Cell 179, 1393-1408.e1316. doi: 10.1016/j.cell.2019.10.025.

Jean, A., Trouillet, A.C., Andrianarivelo, N.A., Mhaouty-Kodja, S., and Hardin-Pouzet, H. (2017). Phospho-ERK and sex steroids in the mPOA: involvement in male mouse sexual behaviour. J Endocrinol 233, 257–267. doi: 10.1530/joe-17-0025.

Johnston, P., and Davidson, J.M. (1972). Intracerebral androgens and sexual behavior in the male rat. Horm Behav 3, 345–357. doi: https://doi.org/10.1016/0018-506X(72)90024-4.

Leranth, C., Petnehazy, O., and MacLusky, N.J. (2003). Gonadal hormones affect spine synaptic density in the CA1 hippocampal subfield of male rats. J Neurosci 23, 1588–1592. doi: 10.1523/jneurosci.23-05-01588.2003.

Lonze, B.E., and Ginty, D.D. (2002). Function and regulation of CREB family transcription factors in the nervous system. Neuron 35, 605–623. doi: 10.1016/s0896-6273(02)00828-0.

Matuszczyk, J.V., and Larsson, K. (1994). Experience modulates the influence of gonadal hormones on sexual orientation of male rats. Physiol Behav 55, 527–531. doi: 10.1016/0031-9384(94)90112-0.

McDonald, P., Beyer, C., Newton, F., Brien, B., Baker, R., Tan, H.S., et al. (1970). Failure of 5alpha-dihydrotestosterone to initiate sexual behaviour in the castrated male rat. Nature 227, 964–965. doi: 10.1038/227964a0.

McEwen, B.S., and Milner, T.A. (2017). Understanding the broad influence of sex hormones and sex differences in the brain. J Neurosci Res 95, 24–39. doi: 10.1002/jnr.23809.

McGinnis, M.Y., and Dreifuss, R.M. (1989). Evidence for a role of testosterone-androgen receptor interactions in mediating masculine sexual behavior in male rats. Endocrinology 124, 618–626. doi: 10.1210/endo-124-2-618.

McGinnis, M.Y., Montana, R.C., and Lumia, A.R. (2002). Effects of hydroxyflutamide in the medial preoptic area or lateral septum on reproductive behaviors in male rats. Brain Res Bull 59, 227–234. doi: 10.1016/s0361-9230(02)00869-9.

Meisel, R.L., and Luttrell, V.R. (1990). Estradiol increases the dendritic length of ventromedial hypothalamic neurons in female Syrian hamsters. Brain Res Bull 25, 165–168. doi: 10.1016/0361-9230(90)90269-6.

Micevych, P.E., and Mermelstein, P.G. (2008). Membrane estrogen receptors acting through metabotropic glutamate receptors: an emerging mechanism of estrogen action in brain. Mol Neurobiol 38, 66–77. doi: 10.1007/s12035-008-8034-z.

Micevych, P.E., Mermelstein, P.G., and Sinchak, K. (2017). Estradiol membrane-Initiated signaling in the brain mediates reproduction. Trends Neurosci 40, 654–666. doi: 10.1016/j.tins.2017.09.001.

Nguyen, T.V., Yao, M., and Pike, C.J. (2009). Dihydrotestosterone activates CREB signaling in cultured hippocampal neurons. Brain Res 1298, doi: 10.1016/j.brainres.2009.08.066.

Parducz, A., Hajszan, T., Maclusky, N.J., Hoyk, Z., Csakvari, E., Kurunczi, A., et al. (2006). Synaptic remodeling induced by gonadal hormones: neuronal plasticity as a mediator of neuroendocrine and behavioral responses to steroids. Neuroscience 138, 977–985. doi: 10.1016/j.neuroscience.2005.07.008.

Paredes, R.G. (2010). “Chapter 13 - Hormones and Sexual Reward,” in Vitamins & Hormones, ed. G. Litwack. Academic Press, 241–262.

Paredes, R.G., Lopez, M.E., and Baum, M.J. (1998). Testosterone augments neuronal Fos responses to estrous odors throughout the vomeronasal projection pathway of gonadectomized male and female rats. Horm Behav 33, 48–57. doi: 10.1006/hbeh.1998.1435.

Pedram, A., Razandi, M., Sainson, R.C., Kim, J.K., Hughes, C.C., and Levin, E.R. (2007). A conserved mechanism for steroid receptor translocation to the plasma membrane. J Biol Chem 282, 22278–22288. doi: 10.1074/jbc.M611877200.

Peterson, B.M., Mermelstein, P.G., and Meisel, R.L. (2015). Estradiol mediates dendritic spine plasticity in the nucleus accumbens core through activation of mGluR5. Brain Struct Funct 220, 2415–2422. doi: 10.1007/s00429-014-0794-9.

Pitchers, K.K., Di Sebastiano, A.R., and Coolen, L.M. (2016). mGluR5 activation in the nucleus accumbens is not essential for sexual behavior or cross-sensitization of amphetamine responses by sexual experience. Neuropharmacology 107, 122–130. doi: 10.1016/j.neuropharm.2016.03.002.

Putnam, S.K., Sato, S., and Hull, E.M. (2003). Effects of testosterone metabolites on copulation and medial preoptic dopamine release in castrated male rats. Horm Behav 44, 419–426. doi: 10.1016/j.yhbeh.2003.06.006.

Rochefort, N.L., and Konnerth, A. (2012). Dendritic spines: from structure to in vivo function. EMBO Rep 13, 699–708. doi: 10.1038/embor.2012.102.

Santollo, J., and Daniels, D. (2019). Anorexigenic effects of estradiol in the medial preoptic area occur through membrane-associated estrogen receptors and metabotropic glutamate receptors. Horm Behav 107, 20–25. doi: 10.1016/j.yhbeh.2018.11.001.

Sargin, D., Mercaldo, V., Yiu, A.P., Higgs, G., Han, J.H., Frankland, P.W., et al. (2013). CREB regulates spine density of lateral amygdala neurons: implications for memory allocation. Front Behav Neurosci 7. doi: 10.3389/fnbeh.2013.00209.

Silva, N.L., and Boulant, J.A. (1986). Effects of testosterone, estradiol, and temperature on neurons in preoptic tissue slices. Am J Physiol 250, R625–632. doi: 10.1152/ajpregu.1986.250.4.R625.

Simerly, R.B., Chang, C., Muramatsu, M., and Swanson, L.W. (1990). Distribution of androgen and estrogen receptor mRNA-containing cells in the rat brain: an in situ hybridization study. J Comp Neurol 294, 76–95. doi: 10.1002/cne.902940107.

Staffend, N.A., Loftus, C.M., and Meisel, R.L. (2011). Estradiol reduces dendritic spine density in the ventral striatum of female Syrian hamsters. Brain Struct Funct 215, 187–194. doi: 10.1007/s00429-010-0284-7.

Staffend, N.A., and Meisel, R.L. (2011). DiOlistic labeling in fixed brain slices: phenotype, morphology, and dendritic spines. Curr Protoc Neurosci Chapter 2, Unit 2.13. doi: 10.1002/0471142301.ns0213s55.

Tabori, N.E., Stewart, L.S., Znamensky, V., Romeo, R.D., Alves, S.E., McEwen, B.S., et al. (2005). Ultrastructural evidence that androgen receptors are located at extranuclear sites in the rat hippocampal formation. Neuroscience 130, 151–163. doi: 10.1016/j.neuroscience.2004.08.048.

Tobiansky, D.J., Wallin-Miller, K.G., Floresco, S.B., Wood, R.I., and Soma, K.K. (2018). Androgen Regulation of the mesocorticolimbic system and executive function. Front Endocrinol 9, 279. doi: 10.3389/fendo.2018.00279.

Tonn Eisinger, K.R., Gross, K.S., Head, B.P., and Mermelstein, P.G. (2018a). Interactions between estrogen receptors and metabotropic glutamate receptors and their impact on drug addiction in females. Horm Behav 104, 130–137. doi: 10.1016/j.yhbeh.2018.03.001.

Tonn Eisinger, K.R., Larson, E.B., Boulware, M.I., Thomas, M.J., and Mermelstein, P.G. (2018b). Membrane estrogen receptor signaling impacts the reward circuitry of the female brain to influence motivated behaviors. Steroids 133, 53–59. doi: 10.1016/j.steroids.2017.11.013.

Vasudevan, N., Kow, L.M., and Pfaff, D. (2005). Integration of steroid hormone initiated membrane action to genomic function in the brain. Steroids 70, 388–396. doi: 10.1016/j.steroids.2005.02.007.

Wallin-Miller, K., Li, G., Kelishani, D., and Wood, R.I. (2016). Anabolic-androgenic steroids decrease dendritic spine density in the nucleus accumbens of male rats. Neuroscience 330, 72–78. doi: 10.1016/j.neuroscience.2016.05.045.

Woolley, C.S., Gould, E., Frankfurt, M., and McEwen, B.S. (1990). Naturally occurring fluctuation in dendritic spine density on adult hippocampal pyramidal neurons. J Neurosci 10, 4035–4039. doi: 10.1523/jneurosci.10-12-04035.1990.

Woolley, C.S., and McEwen, B.S. (1992). Estradiol mediates fluctuation in hippocampal synapse density during the estrous cycle in the adult rat. J Neurosci 12, 2549–2554. doi: 10.1523/jneurosci.12-07-02549.1992.

Zancan, M., Dall’Oglio, A., Quagliotto, E., and Rasia-Filho, A.A. (2017). Castration alters the number and structure of dendritic spines in the male posterodorsal medial amygdala. Eur J Neurosci 45, 572–580. doi: 10.1111/ejn.13460.

